# Human Movement Science in The Wild: Can Current Deep-Learning Based Pose Estimation Free Us from The Lab?

**DOI:** 10.1101/2021.04.22.440909

**Authors:** Laurie Needham, Murray Evans, Darren P. Cosker, Logan Wade, Polly M. McGuigan, James L. Bilzon, Steffi L. Colyer

## Abstract

Human movement researchers are often restricted to laboratory environments and data capture techniques that are time and/or resource intensive. Markerless pose estimation algorithms show great potential to facilitate large scale movement studies ‘in the wild’, i.e., outside of the constraints imposed by marker-based motion capture. However, the accuracy of such algorithms has not yet been fully evaluated. We computed 3D joint centre locations using several deep-learning based pose estimation methods (OpenPose, AlphaPose, DeepLabCut) and compared to marker-based motion capture. Participants performed walking, running and jumping activities while marker-based motion capture data and multi-camera high speed images (200 Hz) were captured. The pose estimation algorithms were applied to 2D image data and 3D joint centre locations were reconstructed. Pose estimation derived joint centres demonstrated systematic differences at the hip and knee (~30 − 50 mm), most likely due to mislabeling of ground truth data in the training datasets. Where systematic differences were lower, e.g., the ankle, differences of 1 − 15 mm were observed depending on the activity. Markerless motion capture represents a highly promising emerging technology that could free movement scientists from laboratory environments. We provide recommendations relating to domain specific datasets and benchmarks, which will be vital to realising this goal.

## INTRODUCTION

The measurement and study of human movement is a fundamental part of many science and medicine disciplines but accurately measuring human motion is an extremely challenging task, even in highly controlled laboratory environments. Examples include, motor rehabilitation scientists and clinicians who quantify movement to inform rehabilitation design^1,2^ and evaluate the effects of disease and treatments^3^. Neuroscientists study brain-movement interaction and motor learning^4^. Similarly, psychologists examine human motor development^5^, human motor behavior^6^ and the effects of psychological disorders on movement^7^. Sports and exercise physiologists and biomechanists examine the metabolic costs of human movement^8^, sports techniques^9,10^, injury mechanisms^11,12^ and equipment design^13^. Finally, engineers quantify movement for prosthetics^14^, exoskeleton^15^ and rehabilitation robotics design^16^. The scope and breadth of these examples demonstrate the substantial impact that that human movement research makes to science and medicine^17^.

The movement sciences described above traditionally rely on vision-based tools using either regular image or video data and manually annotating points of interest or marker-based motion capture systems. Manual annotation of video data is low in cost, easy to deploy in both laboratory and real-world settings and is relatively unobtrusive^10^. However, manual annotation is highly time-consuming and is liable to subjective error^10^, which ultimately limits how much data can be processed and the quality of subsequent analyses. Automated marker-based motion capture systems are now commonplace in laboratory environments providing marker tracking with sub-millimeter accuracy^18^ and greatly reducing the processing time when compared to manual data annotation. However, use of marker-based systems is limited to capturing repetitive, non-representative tasks performed in small, highly controlled environments such as laboratories^17^. The requirement to wear markers may also alter natural movement patterns^10^ and the placement of markers is subject to inter-session and inter-tester variability^19^. Furthermore, marker placements often do not correspond directly to the true anatomical joint centres they are representing^20^, and soft tissue artefact can add further measurement error^21,22^. These limitations of marker-based motion capture systems have been well studied in relation to “gold standard” methods such as bi-planar videoradiography, where marker-based errors up to 30 mm have been reported for lower limb joints centre locations^21,22^. However, marker-based motion capture remains the current de-facto standard for quantifying human movement within laboratories as it is more accessible in terms of cost, flexible in terms of capture volume size and safer, than x-ray based criterion methods.

Pose estimation is a general computer vision problem where the aim is to detect the position and orientation of an object without the placement of markers. Specifically, this process involves detecting a sparse set of key-points that describe the object’s pose. In human pose estimation, joint centres such as the hip, knee or ankle, are estimated in order to reason the position and orientation of the body. This process represents a challenging problem as the algorithm should be invariant to changes in scale, perspective, lighting and even partial occlusion of a body part. The development of large, high-quality data sets (e.g., COCO Keypoint Detection Challenge, MPII Human Pose Dataset and VGG Pose Dataset) has allowed pose estimation research to develop rapidly using supervised deep-learning methods. Typically, some implementation of a convolutional neural network (CNN) is used to learn features associated with each key-point in the training dataset. At deployment, a forward pass of an image returns a 2D confidence map of key point locations (e.g., hip or elbow joint centres) in relation to that image alone. The accuracy of these CNN based methods is typically evaluated against hand-labelled ground truth data which are undoubtably subject to human error^10,17^ and not necessarily a true gold standard measure.

The application of pose estimation algorithms represents an exciting development for movement scientists with the promise of freeing research designs from highly constrained laboratory-based analyses^17^ and allowing for data capture ‘in the wild’. The prospect of using low-cost imaging systems to unobtrusively capture large amounts of data in ecologically valid settings (e.g., in clinics, homes or outdoors) opens new avenues of research with larger and more varied samples, reduced bias and ultimately datasets that better represent the phenomena being studied. But despite the potential that pose estimation presents for the study of human movement, little is known about how accurately such methods can detect human joint centres, which are a fundamental requirement in almost all human motion analysis techniques. Furthermore, to use such an approach within human movement research, considerations must also be made regarding the 3D reconstruction of 2D key-points from multiple cameras and for ensuring robust temporal continuity between frames, e.g., ensuring that key-points are associated or tracked consistently and robustly as a function of time. All of these data processing requirements are non-trivial and require substantial expertise across multiple disciplines (e.g., biomechanics, signal process, computer vision) to produce accurate and reliable markerless 3D representations of human movement.

Human pose estimation is a rapidly developing field of computer science research. Every year there are many releases of new pose estimation algorithms as researchers compete in challenges such as the COCO Keypoint Detection Challenge. While such challenges are designed to accelerate the advancement of pose estimation technology, it is unclear whether these advancements actually benefit pose estimation technology when applied to specific domains such as human movement sciences. It is unfeasible to evaluate every pose estimation algorithm for use in human movement sciences and as such, in this research, we focus upon three popular systems; OpenPose^23^, AlphaPose^24^ and DeepLabCut^25^. OpenPose represents a bottom-up approach to multi-person pose estimation as it simultaneously detects every instance of a given body part while also associating each body part to a person via a set of 2D vector fields that encode the location and orientation of limbs in the image^23^. The OpenPose package was selected here as it is easy to install and use as well as being one of the only systems to provide a foot detection at the time of writing. As such OpenPose is a popular system and has started to receive some attention in movement sciences research^10,26–28^. AlphaPose employs a top-down approach to multi-person pose estimation, first detecting individual people within an image before applying pose-estimation to detect key points of each detected person^24^. AlphaPose was selected as it represents a different approach to pose estimation inference (topdown) and reported improved performance over OpenPose on MPII and COCO dataset benchmarks^24^. Finally, DeepLabCut leverages the DeeperCut pose estimation algorithm^29^, data augmentation and transfer learning to allow researchers to re-train and specialise a CNN to detect user-specified key-points. DeepLabCut provides a set of tools to implement activelearning-based network refinement^30^ and was selected as to date, it has by far had the greatest impact upon movement sciences research^17^.

As with any emerging technology, validation against an established gold-standard is an important step to help researchers understand both the strengths and weaknesses of a system. Such a process permits researchers to make informed decisions about whether emerging technologies provide a suitable tool to help answer their research questions. To date, AlphaPose validation have been limited to post-training model validation, where the final model is validated using human labeled ground truth images provided within the MPII and COCO datasets. However, crowd-sourced datasets are unlikely to have been labeled with the underlying anatomical structures in mind which may reduce the quality of the ground-truth data^31^. DeepLabCut models are also evaluated post-training, however, as training data are typically labelled by researchers with specific domain knowledge, this data and thus the training validation could be of a higher quality than using generic crowd-sourced datasets alone. There remains, however, a need to validate both systems against marker-based motion capture using fully-synchronised, high-speed imaging systems. Two pilot studies have attempted to evaluate the accuracy of OpenPose against marker-based motion-capture using a very small sample size (n=2). Zago et al.,^27^ evaluated 3D joint centre locations derived from a stereo-vision system and OpenPose for walking activities. Joint location differences of between 20 mm and 60 mm were reported. However, unequal sampling frequencies (30 Hz for image data, 100 Hz for marker-based data) and an absence of synchronisation hardware may have contributed to an unknown portion of the reported differences. Nakano et al.,^28^ tested a multi-camera based OpenPose system against marker-based motion capture and reported that 80% of joint centre differences were less than 30 mm. Larger differences were attributed to key-point detection failures at the 2D pose detection stage. Again, there was a lack of synchronisation hardware, low and uneven sampling rates and no information detailing how the two systems Euclidean spaces were aligned making it difficult to know what proportion of the reported differences can be attributed to OpenPose.

There is a need, therefore, to concurrently and robustly evaluate open-source pose estimation algorithms against marker-based motion capture for a range of fundamental human movements. Evaluating using a full-body biomechanical six degrees of freedom model will allow movement scientists to better understand the strengths and weakness of such methods in their current form. Furthermore, testing multiple algorithms will allow for the consistent application of tracking, 3D-fusion and system synchronisation methods, which has so far been absent from the literature to date. The aim of this study was to assess the ability of CNN based pose estimation algorithms to accurately reconstruct the location of joint centres during fundamental human movements. The purpose of this study was to provide human movement scientists with robust evaluation data to make more informed choices about the application of these methods within their own research or practice.

## METHODS

Fifteen healthy participants (7 males [1.82 ± 0.11 m, 85.7 ± 11.1 kg], 8 females [1.65 ± 0.08 m, 63.2 ± 6.0 kg]) provided written informed consent. During a single testing session each participant performed ten walking trials, ten running trials (both at self-selected pace) and ten counter-movement jumps in a randomised order while wearing a full body marker-set. Movement trials were captured concurrently using two motion capture systems. Evaluation data were captured using a 15-camera marker-based motion capture system (Oqus, Qualysis AB, Gothenburg, Sweden) while additional image data were captured using a custom 9-camera computer vision system (JAI sp5000c, JAI ltd, Denmark) (Figure 1). Motion capture systems were time-synchronised by means of a periodic TTL-pulse generated by the custom system’s master frame grabber to achieve a frame locked sampling frequency of 200 Hz in both systems. Additionally, a stopping trigger signal for both systems was generated by the master frame grabber. This ensured that not only did both camera systems stop recording at the same time but that frames were captured by all cameras in unison. To further ensure that synchronisations did not drift, two visible LEDs and one infra-red LED were placed in the capture volume in view of both system’s cameras. The LEDs were programmed to flash in sequence and could later be used to ensure that frame alignment between systems was as expected.

**Figure 1.**
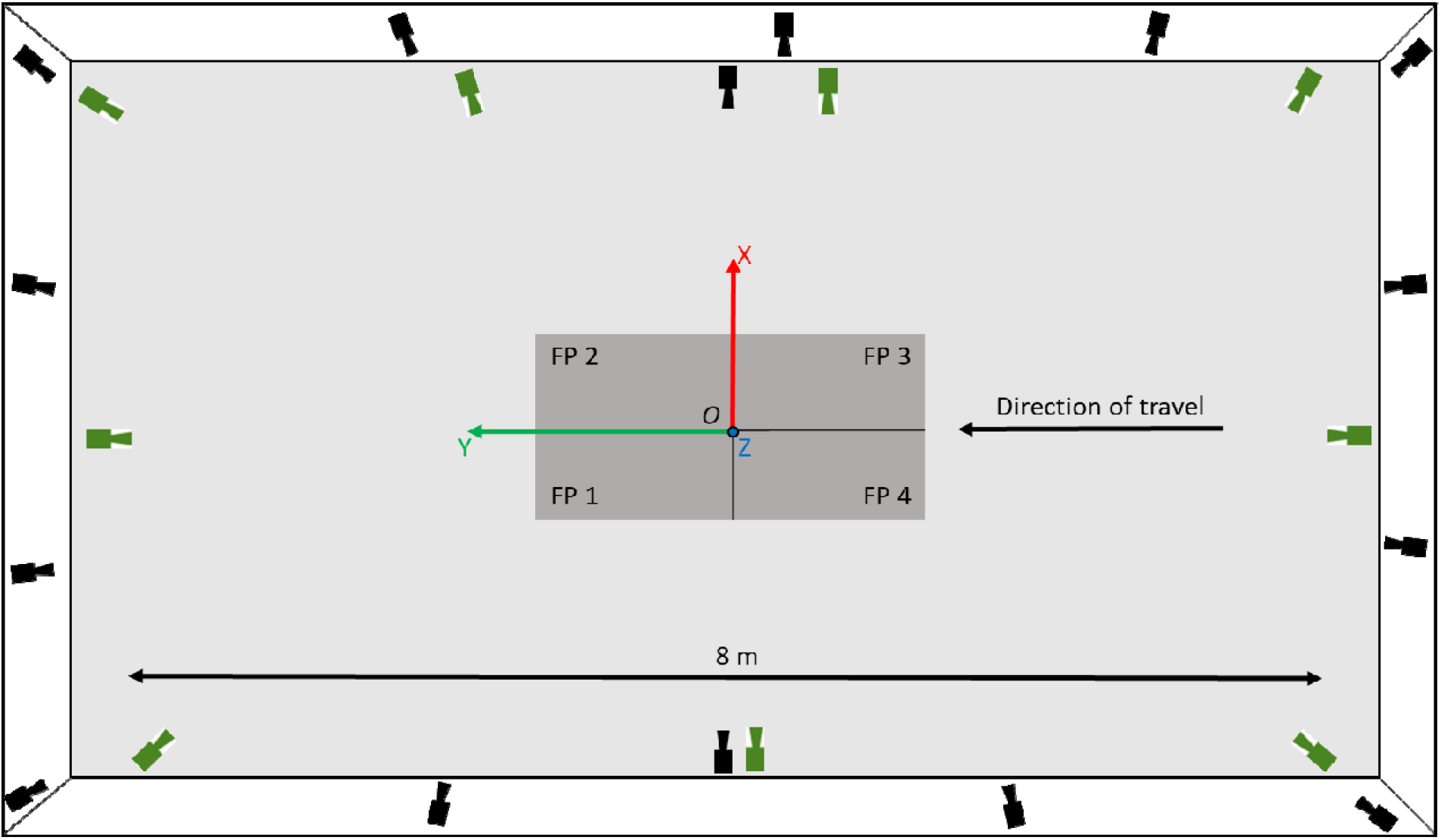
Birds-eye view detailing the layout of the capture volume. Four force plates (FP) are shown in the centre in dark grey. Black cameras depict Qualysis camera locations, green cameras depict JAI machine vision camera locations. The origin of a right-handed coordinate system was set in the centre of the force plates at position *O* with the positive z-axis set normal to the X-Y floor plane.

The Qualysis system was calibrated as per the manufacturer’s specifications. The custom camera system used observations of a binary dot matrix to initialise each camera’s intrinsic parameters^32^, then extrinsic parameters were initialised from pairs of cameras with shared dot matrix observations. A global optimisation was performed using Sparse Bundle Adjustment^33^ to determine the final intrinsic and extrinsic parameters. A right-handed coordinate system was defined for both systems by placing a Qualysis L-Frame in the centre of the capture volume. To refine the alignment of each system’s Euclidean space, a single marker was moved randomly through the capture volume and tracked by both systems. This marker data provided points with which the spatial alignment could be optimised in a least-squares sense. To assess the reconstruction accuracy of both systems a wand was moved through the capture volume and tracked by both systems before the mean (± SD) resultant vector magnitude was computed and compared to the known dimensions of the wand.

To capture criterion data, a full body marker set comprising of 44 individual markers and eight clusters were attached to each participant to create a full body six degrees of freedom (6DoF) model (bilateral feet, shanks and thighs, pelvis and thorax, upper and lower arms, and hands) (Supplementary Materials - Figure A1). Following labelling and gap filling of trajectories (Qualysis Track Manager v2019.3, Qualysis, Gothenburg, Sweden) data were exported to Visual 3D (v6, C-Motion Inc, Germantown, USA) where raw trajectories were low-pass filtered (Butterworth 4^th^ order, cut-off 12 Hz) and a 6DoF model was computed. The marker-based model’s joint centres were computed as the point 50% between the medial and lateral marker for all joints except the hip joint centre which was computed using the regression equations reported by Bell et al^34^.

Multi-camera image data were processed using OpenPose (v1.6.0^23^), AlphaPose (v0.3.0^24^) and DeepLabCut’s pre-trained human pose model (v2.1.7^25^). OpenPose returned a 25-point body model, AlphaPose returned an 18-point body model and DeepLabCut returned a 15-point body model (Supplementary Materials - Figure A2 & Table A1). Temporal frame alignment between marker and markerless systems were confirmed and refined using the flashing LEDs, where required. Tracking and 3D fusion of 2D markerless data were achieved using the method described by^35^. This process first performs occupancy map based cross camera person matching before performing outlier detection and 3D joint reconstruction. 3D joint centre reconstructions were filtered using a bi-directional Kalman filter^36^ before being written to C3D file format. Example videos are provided in the supplementary materials.

For walking and running trials, touch-down (TD) and toe-off (TO) events were computed using Visual 3D’s ‘automatic_gait_events’ function^37^ which was applied to marker-based data before each step cycle was registered to 101 points from TD to the next corresponding TD. As marker and markerless data were temporally synchronised, events derived from marker-based data could be used for markerless data too, thus ensuring event timing consistency between methods. For each trial, an average of four and six complete step cycles were captured for walking and running, respectively. Jumping trials were registered to 101 points from first movement to stabilisation. Where first movement was defined as the point that the vertical force dropped below body weight for 20 consecutive frames and stabilisation was defined as the point that vertical force remained within 3 standard deviations of bodyweight.

Differences in joint centre trajectories between the marker-based and markerless joint centres were determined by computing the 3D Euclidean distance at each time point. Additionally, the signed differences between trajectories were computed along each global coordinate system axis (X-axis: anterior-posterior, Y-axis: medial-lateral, Z-axis: superior-inferior). Agreement between methods was evaluated using Bland-Altman analysis and linear regression models. Normality was tested using a Shapiro-Wilk test. Bland-Altman analysis permits the delineation of systematic (bias) and random (standard deviation of bias) difference between measures with 95% limits of agreement (LoA)^38^. Where data were not found to be normally distributed, non-parametric LoA were computed using the 5^th^ and 95^th^ percentiles^39^. Additionally, we computed linear regression models which provide reliable and sensitive means to compare between biomechanical waveforms^40^. The coefficient of determination (R^2^) indicates the strength of the linear relationship between the two measures while the intercept indicates the shift or offset and the gradient describes the variation of one waveform relative to another^40^.

## RESULTS

Example joint centre trajectories derived from each motion capture method for a single participant during walking, running and jumping are provided in Figure 2. Further examples for other joint centres are provided in the supplementary materials (Figure A3, A4 & A5). For each pose estimation method, mean ± SD time-series differences for the hip joint centre, when compared to marker-based motion capture, are shown in Figure 3. Further joint centre differences are provided in the supplementary materials (Figures A6, A7 & A8).

**Figure 2.**
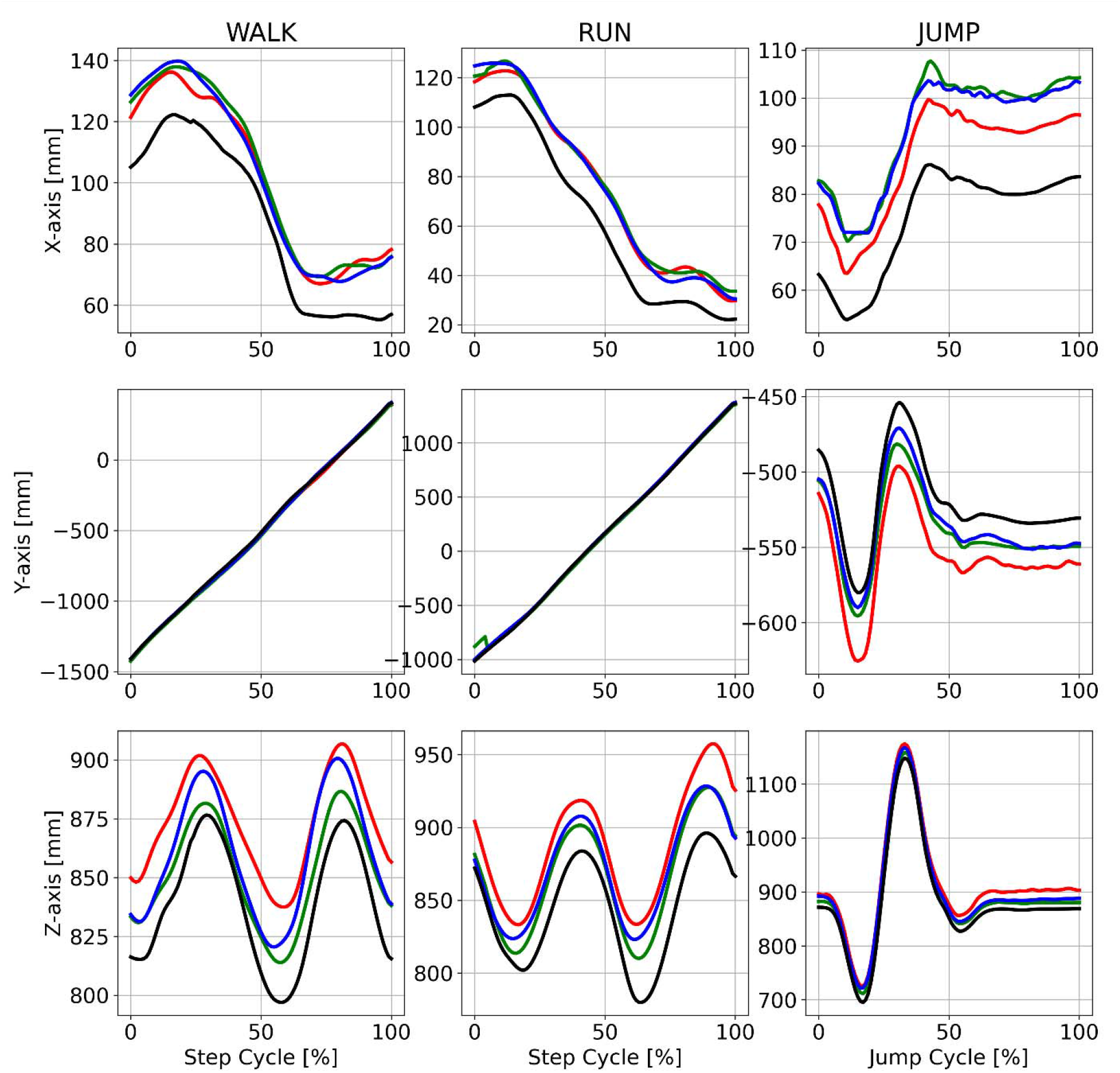
Example mean right hip joint centre trajectories for a single participant (P10) during walking (left), running (centre) and jumping (right) in the global coordinate system axes. Marker based trajectories (black), OpenPose (green), AlphaPose (blue) and DeepLabCut (red). X-axis = medial-lateral. Y-axis = anterior-posterior. Z-axis = superior-inferior.

**Figure 3.**
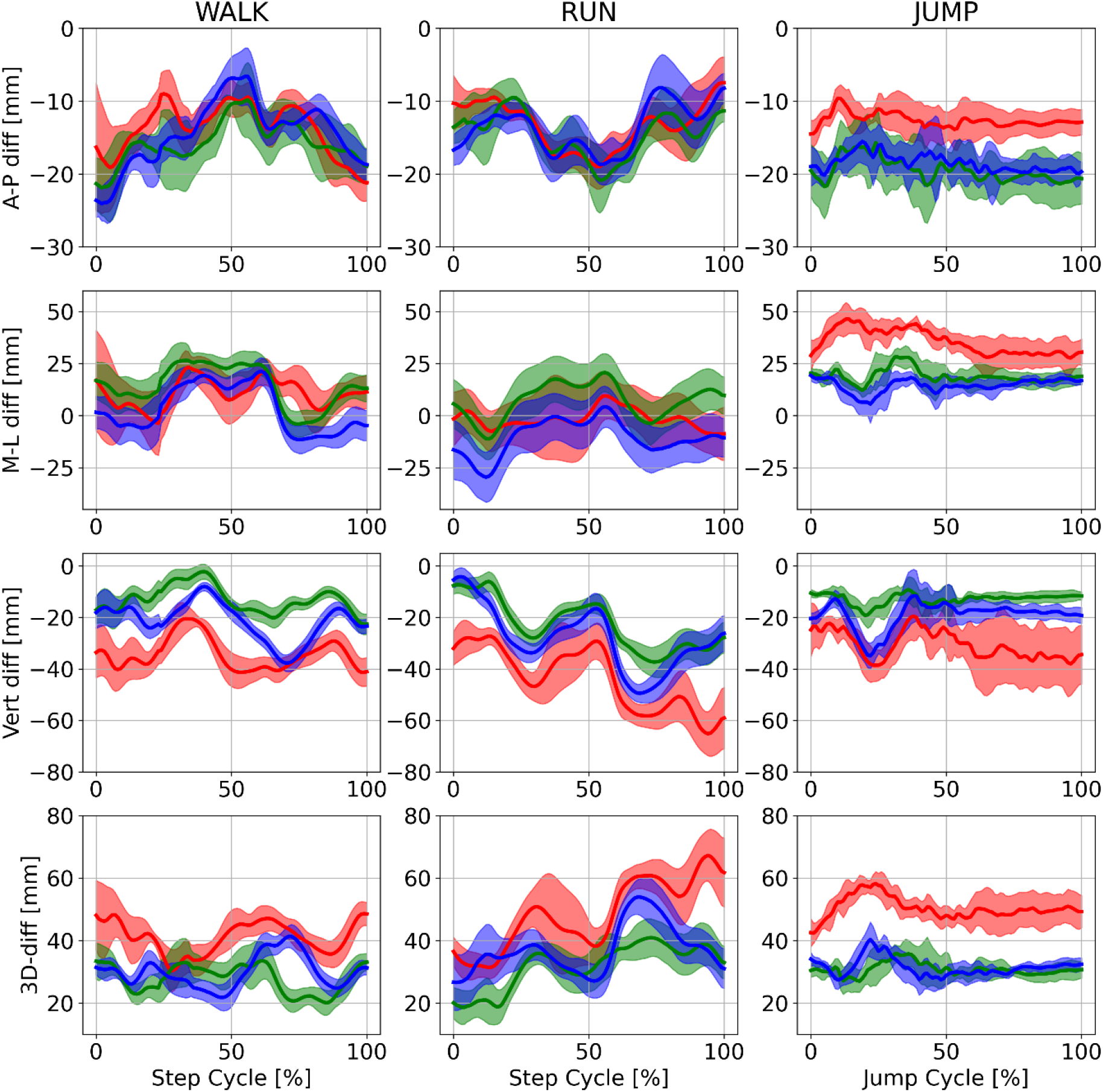
Example mean (± SD) differences between marker based and markerless trajectories for the right hip joint centre trajectories of a single participant (P10) during walking (left), running (centre) and jumping (right). Marker based trajectories (black), OpenPose (green), AlphaPose (blue) and DeepLabCut (red). Row 1 = anterior-posterior differences. Row 2 = medial-lateral. Row 3 = superior–inferior differences. Row 4 = 3D Euclidean differences.

For all pose estimation methods and all activities, the lowest systematic differences were observed at the ankle joint centre (Table 1) with random error and LoA also smallest at this joint. The hip and knee joint centres displayed the largest systematic differences for all pose estimation methods (Table 1 and Figure 4) with knee demonstrating higher random error and LoA during walking and running. The knee also presented higher random error and LoA during jumping. Shoulder joint centre mean differences were typically observed to be larger than those at the ankle but lower than the hip or knee with random error and LoA following the same trend. OpenPose difference distributions for each joint centre are provided in Figure 4 with further examples for AlphaPose and DeepLabCut provided in the supplementary materials (Figure A11 & A12).

**Table 1.** Bland-Altman analysis results of 3D Euclidean differences during walking, running and jumping. OP = OpenPose, AP = AlphaPose, DLC = DeepLabCut.

| | | Mean Difference (Bias) (mm) | | | Random Error (± SD of Bias) | | | 95% LoA (Lower - Upper Limit [mm]) | | | | | |
| --- | --- | --- | --- | --- | --- | --- | --- | --- | --- | --- | --- | --- | --- |
|  |  | OP | AP | DLC | OP | AP | DLC | OP |  | AP |  | DLC |  |
| WALK | Shoulder | 29 | 30 | 24 | 8 | 7 | 8 | 18 | 43 | 21 | 42 | 14 | 37 |
|  | Hip | 34 | 31 | 43 | 12 | 9 | 14 | 14 | 53 | 15 | 45 | 21 | 68 |
|  | Knee | 30 | 31 | 42 | 11 | 15 | 26 | 15 | 51 | 13 | 61 | 17 | 84 |
|  | Ankle | 16 | 19 | 30 | 8 | 14 | 39 | 6 | 32 | 6 | 50 | 8 | 65 |
| RUN | Shoulder | 27 | 31 | 29 | 9 | 11 | 13 | 13 | 43 | 15 | 50 | 10 | 51 |
|  | Hip | 29 | 32 | 45 | 9 | 10 | 21 | 14 | 45 | 16 | 50 | 18 | 78 |
|  | Knee | 41 | 48 | 58 | 17 | 26 | 36 | 19 | 74 | 17 | 97 | 22 | 116 |
|  | Ankle | 23 | 36 | 52 | 14 | 24 | 68 | 6 | 50 | 9 | 79 | 11 | 121 |
| JUMP | Shoulder | 32 | 34 | 33 | 11 | 12 | 11 | 16 | 53 | 18 | 56 | 19 | 53 |
|  | Hip | 36 | 36 | 53 | 11 | 10 | 15 | 18 | 54 | 20 | 51 | 29 | 78 |
|  | Knee | 29 | 27 | 35 | 8 | 9 | 9 | 15 | 44 | 14 | 43 | 20 | 49 |
|  | Ankle | 14 | 14 | 15 | 6 | 7 | 8 | 6 | 25 | 4 | 27 | 6 | 29 |

**Figure 4.**
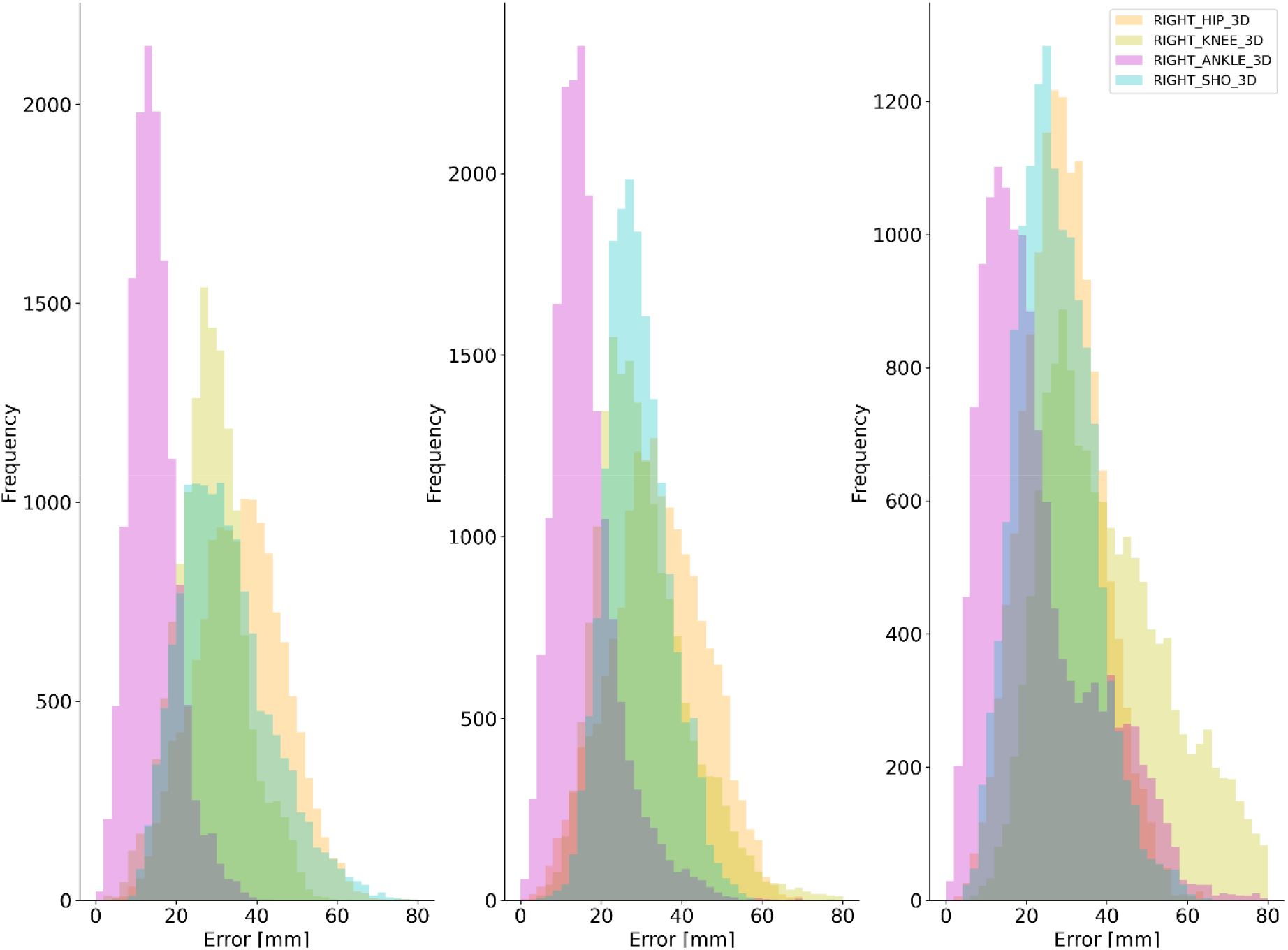
Example difference distributions for OpenPose at the right shoulder, hip, knee and ankle joint centres for jumping (left), walking (centre) and running (right).

**Figure 5.**
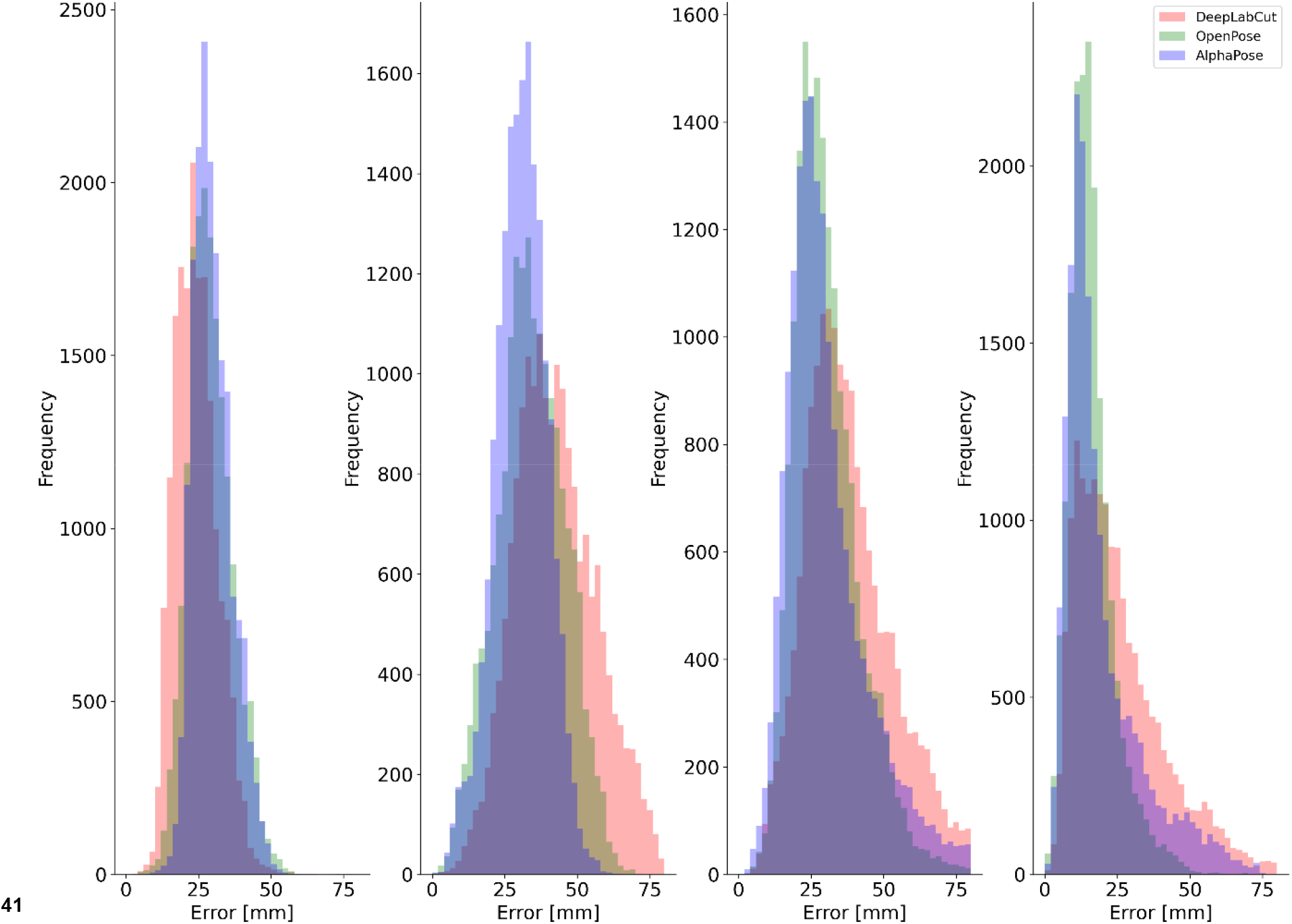
Difference distributions during walking for each pose estimation method at the right shoulder (left), hip (centre-left), knee (centre-right) and ankle (right) joint centres.

Differences in pose estimation performance were observed between walking, running and jumping. For walking and running, the smallest systematic and random differences were observed for OpenPose and largest for DeepLabCut. However, during jumping OpenPose and AlphaPose mean differences and random errors were comparable between markerless methods (within ~ 1 mm of each other) (Figure A10). While the largest systematic differences, random errors and LoA were generally observed for DeepLabCut, the results for ankle and shoulder during jumping were comparable to those observed for OpenPose and AlphaPose. Further visual examples are provided as videos in the supplementary materials.

## DISCUSSION

This is the first study to robustly assess the ability of multiple CNN based pose estimation algorithms (OpenPose^23^, AlphaPose^24^ and DeepLabCut^25^) to accurately reconstruct the location of joint centres during fundamental human movements. Markerless pose estimation algorithms show great potential to facilitate large scale movement studies in a range of environments and the results of this study align with others^28^ in demonstrating that the resulting 3D skeletons *can* under favorable conditions, produce visually impressive outcomes using several different pose estimation methods. However, for all three pose estimation methods during all activities (walking, running and jumping), there was clear evidence of both systematic and random differences when compared to marker-based motion capture (Table 1, A2 A3 & A4), which suggests that such technology may require further development before its performance surpasses the current de-facto methods (marker-based motion capture) in human movement sciences.

The largest differences between pose estimation derived 3D joint centres were observed at the hip, where mean differences ranged between 29 mm (OpenPose during running) and 53 mm (DeepLabCut during running). Bland-Altman analysis revealed that these differences were largely systematic in nature with all pose estimation methods placing the hip joint centres in a more lateral and inferior position than those derived from markers placed on the pelvis (Table A2 A4 & A6). Additionally, systematic differences were observed at the knee joint, most notably in the vertical axis where all pose estimation methods placed the joint centre between ~20 mm – 30 mm below the marker derived joint centre. Deep learning-based pose estimation methods are typically trained using supervised learning techniques where annotated images provide training examples of the desired joint centre locations. Systematic differences in joint centre locations such as those observed here are likely a result of large-scale mislabeling of hip and joint centre locations in the datasets used to train each deep learning model. At best, we can expect that supervised deep learning models will be able to perform as well as but not better than the data with which they have been trained^42^. Consequently, it is unrealistic to expect a pose estimation model trained on open access data sets such as COCO^43^ or MPII^44^ to estimate the location of anatomically accurate joint centres when as this study suggests, such joint centres are not accurately labelled in the training data. There is a need, therefore, for the development of large scale, open access pose estimation data sets that have been labelled by individuals with detailed knowledge of human anatomy. Although systematic and random errors were largest for DeepLabCut, the software provides functionality for leveraging transfer learning, which helps specialise pre-trained networks using small amounts of additional training data. As such, we would expect to see improved results for DeepLabCut if additional data and training time is leveraged on top of the DeepLabCut pre-trained human pose model that was evaluated in this study. Indeed, previous work utilising re-trained DeepLabCut has demonstrated promising 2D sagittal plane results during underwater running with mean differences of approximately 10 mm^45^.

The smallest differences were observed for the ankle joint, which demonstrated considerably lower systematic differences and random errors when compared to the hip and knee joint centres. The results observed for the ankle joint centre, which is perhaps easier to identify and label than the hip and knee demonstrate the potential that, given (conceivably more) anatomically accurate training data, pose estimation methods could achieve for all joint centres. Indeed, mean differences as low as 2 ± 7 mm (Table A4 & A5) were observed using OpenPose during jumping and represent values that are well within the known error ranges (25 – 33 mm) of marker-based motion capture techniques^21,22^. Once all joints can be estimated to this level of accuracy the focus of attention instead turns to between session repeatability^46^, which can be problematic for marker-based systems when markers are not placed in identical locations between sessions^47^.

Evidence of activity specific pose estimation performance was observed, with the largest mean differences and random error measured during running and the smallest mean differences and random error measured during jumping. Lower pose estimation performance during running may be attributed to the greater range of observed limb configurations and segment velocities when compared to jumping and aligns with previous evidence that pose estimation performance is highly task specific^35^ - an important consideration for researchers. Furthermore, greater limb velocities may introduce image noise in the form of motion blur, making it harder for pose estimation methods to detect image features that represent a given joint centre. Indeed, it is unlikely that current datasets include training examples with any degree of motion blur, a factor which should be considered during the development of future datasets. In this study, we minimised noise from motion blur by utilising high frame rates (200 Hz), low exposure times and high-quality studio lighting. Yet, there exists a trade-off between capturing the highest quality images and using pose estimation models that can handle less than perfect images, which researchers should carefully consider as part of their research design.

Performance of OpenPose and AlphaPose across activities and joints was comparable to one another for systematic differences (between method differences ~1 – 5 mm) and random errors (between method differences ~1 – 3 mm) while larger systematic and random errors were observed for DeepLabCut. Despite using distinctly different network architectures and approaches to pose estimation, the comparable performances of OpenPose and AlphaPose were surprising. AlphaPose has previously demonstrated superior performance on computer vision benchmarks such as COCO and MPII. However, as the results of this study demonstrate, pose estimation performance should not be solely based upon these computer vision benchmarks but additionally, domain specific benchmarks using appropriate ground truth information are required^17^ to enhance their development and ability to generalise effectively in real-world applications. DeepLabCut did not perform as well as OpenPose and AlphaPose. DeepLabCut uses DeeperCut^29^ to perform pose estimation, which is the oldest method and lowest scoring (on MPII benchmark) method tested in this study and may partially explain the lower performance. An additional source of error that was unique to DeepLabCut related to single and multi-person detection capabilities. The pre-trained human pose model used in this study only returned pose information for a single person. When multiple people were in the field of view, there was a tendency for joint centres to jump between the study participant in the foreground and people in the background (supplementary video – DLC_2Dv3D). Such outliers were largely removed during 3D fusion, however it is inevitable that this issue will have contributed at times to the overall differences that were observed for DeepLabCut. It is worth noting, however, that DeepLabCut provides comprehensive tools for labelling additional data and re-training their pose estimation models. As mentioned previously, by leveraging DeepLabCut’s transfer learning capabilities we would expect to see substantially improved results on those presented in this study. However, the aim of this study was to assess the current capabilities of pre-trained pose estimation models. For all pose estimation methods, our results include larger errors caused by issues such as false positive detections, tracking failures and erroneous switching of limbs. While previous research has suggested manually correcting errors^28^, we did not feel that this approach provided the full picture regarding pose estimation. Rather it is important for those applying pose estimation methods to be fully aware of the issues discussed in this study that require effective detection and correction during the data processing pipeline.

Aligning with previous studies examining OpenPose vs marker-based motion capture^27,28^, we have shown promising face validity for 3D joint centre locations detected using OpenPose, AlphaPose and DeepLabCut but results were not consistently comparable to marker-based motion capture. It is not possible to state that the results of this study meet the accuracy requirements for all human movement science studies, as such requirements will vary greatly between applications and researchers should consider carefully how appropriate the use of current markerless methods are within the context of their research questions. However, our results for joint centres such as the ankle and shoulder do fall within the known range of error for marker-based motion capture^21^ demonstrating that markerless technology has a promising future in human movement science research. There are three key areas to consider for future development of markerless motion capture. Firstly, the need for domain specific datasets which contain anatomically accurate labels and contain representative images for the activity of interest. Additionally, ensuring that each segment has at least three non-colinear keypoints will better facilitate 6DoF pose estimation. Secondly, the need for domain specific benchmarks^17^ that go beyond the current computer vision benchmarks to test pose estimation methods on domain specific ground truth data. For example, in this study we benchmarked against validated marker-based joint centre estimation methods rather than hand labelled joint centre annotations. Without these two considerations (datasets and benchmarks) we will not see significant progress for pose estimation in the movement sciences. Finally, using modelling methods such as inverse kinematics optimisation^48^, which is widely established in many parts of human movement sciences, we would expect that the 3D joint centres derived from pose estimation methods, such as those presented in this study, could be used to provide improved estimates of 3D joint centres and segment kinematics. It is important to reiterate that the evaluation measure used in this study (marker-based motion capture) does not represent the true criterion measure (bi-planar videoradiography) and findings should be considered in this context. However, it is indeed promising that our results often fall within the known error ranges of marker-based motion capture and future developments should only strengthen the capabilities of markerless motion capture to free researchers from the laboratory to perform large scale studies outside of laboratory environments.

## CONCLUSIONS

In this study we demonstrated that OpenPose, AlphaPose and DeepLabCut can be used to detect and reconstruct markerless 3D joint centre locations. When compared to marker-based motion capture, systematic differences were observed at the hip and knee (~30 – 50 mm for all methods), most likely due to mislabeling of ground truth data in the training datasets. Where systematic differences were lower, e.g., at the ankle, differences of 1 – 15 mm ± 10 mm were observed. OpenPose and AlphaPose demonstrated comparable performance to one another and outperformed DeepLabCut. Markerless pose estimation using the methods described in this study do not yet match the performance of marker-based motion capture; however, pose estimation continues to improve at a rapid rate and in line with the suggestions outlined above, could soon open new and less restricted opportunities for researchers wishing to operate outside of the laboratory to capture human movement ‘in the wild’.

## ACKNOWLEDGEMENT

This research was part-funded by CAMERA, the RCUK Centre for the Analysis of Motion, Entertainment Research and Applications, EP/M023281/1 and EP/T014865/1

## CONTRIBUTIONS

The work was conceived by LN, ME, DC, LW, PM, SC. LN, ME & LW collected the data. LN & ME processed the data. LN analysed the data and prepared manuscript and figures. All authors (LN, ME, DC, LW, PM, JB, SC) revised and reviewed the manuscript. Funding was acquired by DC & JB.

## ETHICS DECLARATIONS

The study was conducted according to the guidelines of the Declaration of Helsinki, and approved by the Institutional Review Board (or Ethics Committee) of the University of Bath (EP1819052 25/07/19).

